# Improved memory CD8 T cell response to delayed vaccine boost is associated with a distinct molecular signature

**DOI:** 10.1101/2022.08.04.502772

**Authors:** Ambra Natalini, Sonia Simonetti, Gabriele Favaretto, Lorenzo Lucantonio, Giovanna Peruzzi, Miguel Muñoz-Ruiz, Gavin Kelly, Alessandra Maria Contino, Roberta Sbrocchi, Simone Battella, Stefania Capone, Antonella Folgori, Alfredo Nicosia, Angela Santoni, Adrian C. Hayday, Francesca Di Rosa

**Affiliations:** Institute of Molecular Biology and Pathology, National Research Council of Italy (CNR), Rome 00161, Italy; Department of Molecular Medicine, University of Rome “Sapienza”, Rome 00161, Italy; Center for Life Nano- & Neuro-Science, Fondazione Istituto Italiano di Tecnologia (IIT), Rome 00161, Italy; Immunosurveillance Laboratory, The Francis Crick Institute, London NW11AT, UK; Bioinformatic and Biostatistics Science & Technology Platform, The Francis Crick Institute, London NW11AT, UK; ReiThera S.r.l., Rome 00128, Italy; CEINGE, Naples 80145, Italy; Department of Molecular Medicine and Medical Biotechnology, University of Naples Federico II, Naples 80131, Italy; IRCCS Neuromed, Pozzilli (Isernia) 86077, Italy; Peter Gorer Department of Immunobiology, King’s College London, London SE1 9RT, UK; National Institute for Health Research (NIHR) Biomedical Research Center (BRC), Guy’s and St Thomas’ NHS Foundation Trust and King’s College London, London SE1 9RT, UK

**Keywords:** CD8 T cells, vaccination, prime-boost interval, transcriptomic profile, memory

## Abstract

Although the prime/boost interval can impact vaccine responses, the criteria for deciding its time length are poorly defined. To address this, we examined CD8 T cell responsiveness to boost in a BALB/c mouse model of intramuscular (i.m.) vaccination by priming with HIV-1 gag-encoding Chimpanzee adenovector, and boosting with HIV-1 gag-encoding Modified Vaccinia virus Ankara. We found that boost was more effective at day(d)100 than at d30 post-prime, as evaluated at d45 post-boost by multi-lymphoid organ assessment of gag-specific CD8 T cell frequency, CD62L-expression (as a guide to memory status) and *in vivo* killing. RNA-sequencing of splenic gag-primed CD8 T cells at d100 revealed a quiescent, but highly responsive signature, that trended toward a central memory (CD62L^+^) phenotype. Interestingly, gag-specific CD8 T cell frequency selectively diminished in the blood at d100, relative to the spleen, lymph nodes and bone marrow. These results move forward the rational design of prime/boost intervals.

## Introduction

CD8 T cells are one of the pillars of adaptive immunity. Naïve CD8 T cells are primed in lymph nodes (LNs) and the spleen by mature dendritic cells (DCs), which present short antigen (Ag)-derived peptides in the groove of Major Histocompatibility Complex (MHC) class I (MHC-I) molecules, together with sufficient costimulatory signals and in the context of CD4 T cell help. Thereupon, Ag-responding CD8 T cells proliferate and differentiate as effectors and memory cells. In contrast to naïve T cells that mostly recirculate in blood, spleen and LNs, effector and memory T cells have enhanced capacity to migrate to the bone marrow (BM) and extra-lymphoid tissues (Di Rosa and Gebhardt, 2016). Upon recognition of Ag-MHC-I presented by target cells, effector CD8 T cells display cytotoxic activity and/or cytokine production, thus critically contributing to the clearance of Ag-expressing cells. Afterwards most effector CD8 T cells die, while a few memory CD8 T cells remain durable, ready to provide an enhanced secondary response in case of subsequent encounter with the same Ag (Harty and Badovinac, 2008).

Although CD8 T cell responses are critical for protection against viral infections and tumors (Appay et al., 2008), the development of CD8 T cell-eliciting vaccines proved challenging for many years. The obstacles have now been reduced by some effective platforms, including those based on adenoviral vectors (Bolinger et al., 2015; Capone et al., 2010; Majhen et al., 2014) and on mRNA (Cagigi and Loré, 2021). Nevertheless, there is no clear protocol for how best to induce long-lasting immunity in a predictable manner, e.g., in respect to vaccine dose, route of administration, and time intervals between repeated injections. Solving these issues can be extremely beneficial for immunological understanding and for public health policy, as emphasized by the current COVID-19 pandemic (Krammer, 2020; Lipsitch et al., 2022).

We previously showed that the Chimpanzee adenovector ChAd-gag induced a strong clonal expansion of CD8 T cells against the model antigen Human Immunodeficiency Virus-1 (HIV-1) gag in BALB/c mice (Simonetti et al., 2019). Indeed, after intramuscular (i.m.) injection with ChAd-gag, gag-specific CD8 T cells in S-G_2_/M phases of cell cycle were found not only in LNs and spleen, but also in peripheral blood. Results were similar after boosting with Modified Vaccinia virus Ankara (MVA)-gag (Simonetti et al., 2019). Using this ChAd-gag/MVA-gag model (Colloca et al., 2012; Simonetti et al., 2019), we have now addressed here the impact of the prime/boost time interval on gag-specific CD8 T cell immunity. Our hypothesis was that establishment and regulation of quiescence in primed CD8 T cells after clonal expansion would be closely related to the cells’ responsiveness to boosting. To test this hypothesis, we evaluated in parallel the post-proliferative tail of the primary response in spleen, LNs, BM and blood, and the kinetics of responsiveness to boost. We identified the time-shift from low to high responsiveness, and characterized the molecular profile of highly responsive splenic memory CD8 T cells.

## Results

### Kinetics of gag-specific CD8 T cell expansion and T_CM_-phenotype changes after prime

We exploited our recently developed DNA/Ki-67 flow cytometry assay (Simonetti et al., 2019; Muñoz-Ruiz et al., 2020; Natalini et al., 2021) to evaluate the kinetics of expansion of CD8 T cells specific for the immunodominant gag_197-205_ peptide (gag-specific) in spleen and LNs of ChAd-gag-primed BALB/c mice, up to ∼ 3 months after prime (Fig. 1 and S1). As depicted in Fig. S1A, our strategy of analysis included a DNA-based singlet gate (Step 1), and an unusually “relaxed” FSC-A/SSC-A (FSC-SSC) gate (Step 4, in orange), as opposed to a classical “narrow” gate (Step 4, shown in white for comparison), as previously described (Simonetti et al., 2019; Simonetti et al., 2021; Muñoz-Ruiz et al., 2020; Natalini et al., 2021). Typical spleen gag-specific CD8 T cell profiles are shown in Fig. 1A, and examples of DNA/Ki-67 plots in Fig. 1B, with gating of cells in G_0_, G_1_ and S-G_2_/M phases of cell cycle. The frequency of spleen gag-specific CD8 T cells increased from d10 to d30, and then was maintained at a plateau of ∼0.7% in primed mice, with a background of 0.0-0.1% in untreated control mice (Fig. 1C). Early after priming 83% and 10% of spleen gag-specific cells were on average in G_1_ and in S-G_2_/M respectively, with very similar results at d10 and d14. At d30 and later time points, the percentage of cells in S-G_2_/M dropped to virtually none, while that of cells in G_1_ slowly declined. The G_1_ trend was mirrored by the gradual increase of cells in G_0_, which represented the majority of gag-specific cells from day 35 onwards (Fig. 1D). To track the kinetics of re-entry of spleen gag-specific CD8 T cells into a resting state after clonal expansion, we exploited FSC and SSC changes associated with cell cycle (Simonetti et al., 2019; Muñoz-Ruiz et al., 2020). Thus, we examined in parallel the percentages of gag-specific cells comprised within either the “narrow” FSC-SSC (Fig. 1E) or the FSC-A/-H gate (Fig. 1F). While these gates are normally applied to total spleen cells for lymphocyte and single cell gating, respectively, we exploited them here to follow Ag-primed T cell changes over time. Only a fraction of spleen gag-specific CD8 T cells was captured by any of these two gates in the second week post-prime (Fig. 1E-G), as expected (Simonetti et al., 2019). This inadequacy was more evident in the case of the “narrow” FSC-SSC gate than in that of the FSC-A/-H gate (Fig. 1E-G). One month after priming (d30-d35), the gag-specific cell percentage within the “narrow” FSC-SSC gate raised to ∼88-93%, and that within the FSC-A/-H gate to ∼98-99%, thus resembling the corresponding percentage of not gag-specific cells; there was no further change at d66 and d77 (Fig. 1E-G). Results were similar in draining LNs (Fig. S1B-E). Further analysis demonstrated that only a small percentage of cells in S-G_2_/M was captured by the FSC-A/-H gate, and almost none by the “narrow” FSC-SSC gate in both LNs and spleen (Fig. S1F), in agreement with our previous data (Simonetti et al., 2019).

**Figure 1.**
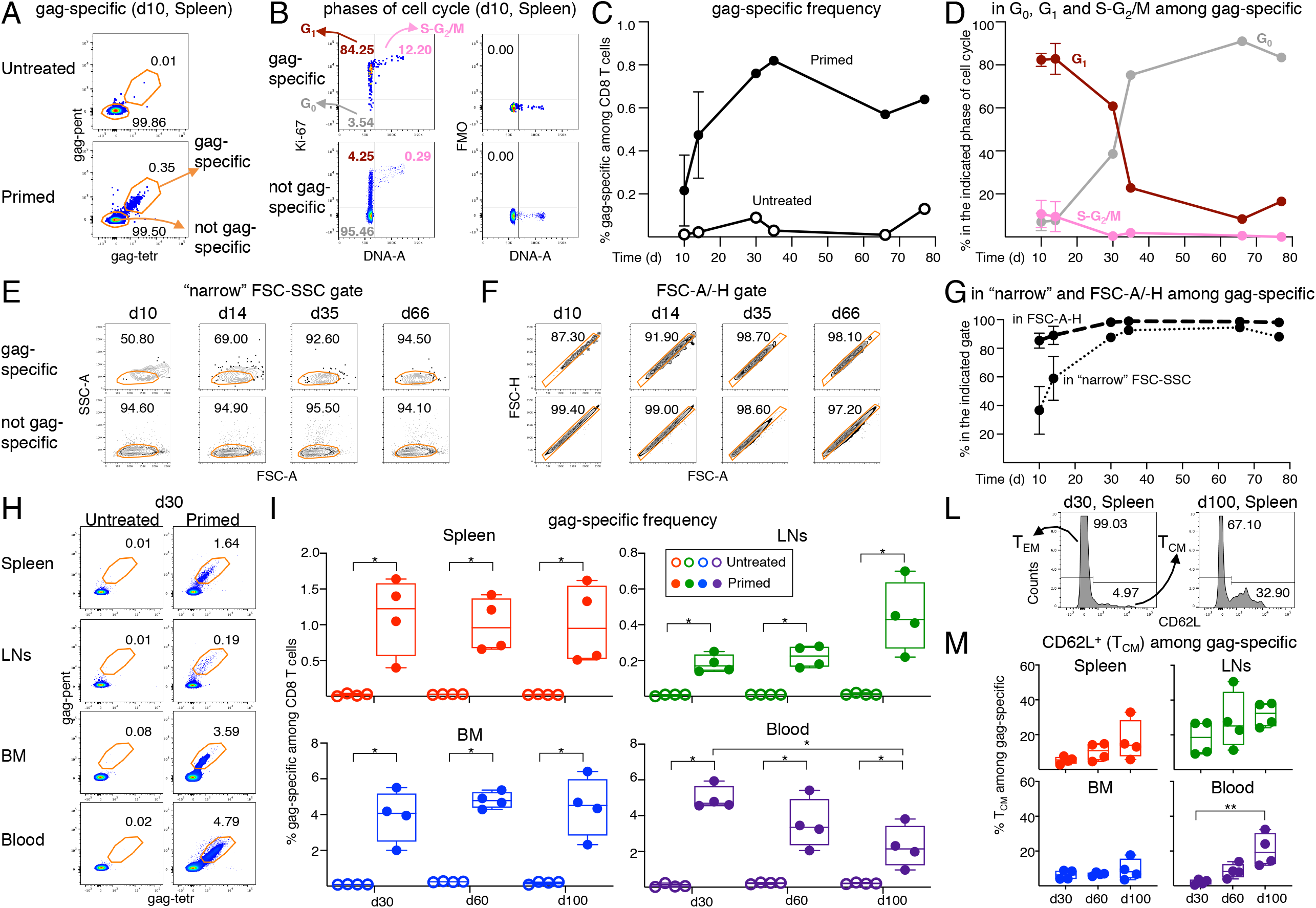
Analysis of frequency, cell cycle and T_CM_/T_EM_-phenotype of gag-specific CD8 T cells from ChAd-gag-primed mice. Female BALB/c mice were primed i.m. in the quadriceps with ChAd-gag (10^7^ vp) at d0 and the kinetics of gag-specific CD8 T cell response was tracked by flow cytometry. **(A-G)**. Spleen cells were analysed by membrane and Ki-67/DNA staining at different days (d) post-prime (gating strategy in fig. S1A). Typical plots showing the percentage of “gag-specific” and “not gag-specific cells” from untreated (top) and primed (bottom) mice at d10 (A). gag-specific cells from primed mice were further analyzed on DNA/Ki-67 plots as follows (B, top left panel): cells in the G_0_ phase of cell cycle were identified as DNA 2n/ ki-67^−^ (bottom left quadrant); cells in G_1_ as DNA 2n/ Ki-67^+^ (upper left quadrant); cells in S-G_2_/M as DNA>2n/ Ki-67^+^ (top right quadrant). As a comparison, not gag-specific cells (B, bottom left panel) are shown. For each DNA/Ki-67 plot, corresponding Ki-67 Fluorescence Minus One (FMO) control plots (B, right panels) are shown. Summary of the kinetics of gag-specific frequency in primed and untreated mice (C), and of cell cycle phases of gag-specific CD8 T cells in primed mice (D). Kinetics of the percentages of gag-specific CD8 T cells in the “narrow” FSC-SSC gate and in the FSC-A/-H gate (examples in E and F, respectively, and summary in G). **(H-M)**. Spleen, lymph nodes (LNs), bone marrow (BM) and blood cells were analysed by membrane and Ki-67 staining at different days (d) post-prime (gating strategy in fig. S1H, Ki-67 results in Fig. 2). Typical plots showing the percentage of gag-specific CD8 T cells at d30 in untreated (left) and primed (right) mice (H) and summary of results at d30, d60 and d100 (I). Examples of CD62L histograms of spleen gag-specific CD8 T cells at d30 and d100, showing the gate used to discriminate between T_CM_ (CD62L^+^) and T _EM_ (CD62L^—^) cells (L). Summary of T_CM_ percentages at d30, d60 and d100 (M). In flow cytometry plots (in A, B, E, F, H, and L) the numbers represent the percentages of cells in the indicated regions. Panels C, D, and G summarize results of 6 independent prime experiments with a total of 84 mice analyzed at the indicated d post-prime. At d10 and d14, symbols represent the mean and bars the Standard Deviations of 5 experiments (each performed with pooled cells from 3 mice per group). At d30, d35, d66 and d77, each symbol represents a pool of 3 mice. Panels I and M summarize results of 4 independent prime experiments with a total of 72 mice. Each symbol represents a pool of 3 mice. Statistical analysis was performed using Mann-Whitney test for comparison between untreated and primed mice (I), and Kruskal-Wallis test with Dunn’s correction for multiple comparison for comparison among primed mice at d30, d60 and d100 (I and M). Statistically significant differences are indicated (* *P* ≤ 0.05; ** *P* ≤ 0.01). This figure includes unpublished data in relation to (Simonetti et al., 2019).

We then used flow cytometry to track at d30, d60 and d100 post-prime the frequency of gag-specific cells in spleen, LNs, BM and blood, and the proportions among them of cells either expressing membrane CD62L, i.e. having a T_CM_ phenotype (Sallusto et al., 2004; Ivetic et al., 2019), or lacking intracellular Ki-67, i.e. being in the quiescent phase G_0_ (Di Rosa et al., 2021). Ki-67 and CD62L analysis were also combined, as explained in the next paragraph. It should be noted that in these experiments we did not stain DNA and relied on the typical FSC-A/-H gate to exclude cell aggregates (Fig. S1G, step 1), since we had observed that gag-specific cells in S-G_2_/M were extremely rare after d30 (Fig. 1D and S1E) and that 98-99% of gag-specific cells were comprised in the FSC-A/-H gate at d30 and onwards (Fig. 1F-G, and S1D). We used a “relaxed” FSC-SSC gate (Fig. S1G, step 4), that was more appropriate than a “narrow” FSC-SSC gate for evaluation of gag-specific cells in G_1_ (see Fig. S1F), even after d30 when G_1_ mostly represented a post-mitotic state (Fig. 1D and S1E). We found that gag-specific frequency at d30 was on average 5.0% in blood, 3.8% in BM, 1.1% in spleen and 0.2% in LNs from primed mice (examples of flow cytometry plots in Fig. 1H, summary of results in Fig. 1I). This frequency remained roughly stable at d60 and d100 in the spleen and BM, with a tendency to increase in the LNs. In contrast, a significant decline was observed in the blood from d30 to d100 (Fig. 1I). At any time point, and in each organ, primed mice had a significantly higher gag-specific frequency than untreated controls, which always displayed a negligible background (Fig. 1H-I). In the primed mice samples, we discriminated between T_CM_ and T_EM_ gag-specific CD8 T cells according to their CD62L expression (Fig. 1L). We found that the proportion of T_CM_ tended to be higher in the LNs, and to increase over time in all organs (Fig. 1M). Notably, there was a significant rise in T_CM_ proportion among gag-specific CD8 T cells in the blood, from ∼2% at d30 to ∼21% at d100 on average (Fig. 1M), reflecting the contemporary changes in lymphoid organs.

### Kinetics of Ki-67 expression loss in gag-specific CD8 T_CM_ and T_EM_ cells after prime

We reasoned that the reported difference in proliferative potential between T_EM_ and T_CM_ cells (Sallusto et al., 2004) might be related to diversity in their kinetics of re-entry into the G_0_ phase after priming. We thus tracked variations in ki-67^−^ (i.e. in G_0_) T_EM_ and T_CM_ cells in the primed mice samples described above. At d30, ki-67^−^cells were on average ∼81% in T_CM_ and ∼68% in T_EM_ gag-specific cells from primed spleen (examples of flow cytometry histograms in Fig. 2A, top). Both percentages increased over time, being on average ∼96% in T_CM_ and ∼94% in T_EM_ at d100 (examples in Fig. 2A, bottom). The marked Ki-67^+^ cell decline in T_EM_ from d30 to d100 was concurrent with a higher representation of T_CM_ cells at d100, as evident in a typical Ki-67/ CD62L plot showing an overlay of T_CM_ (gray) and T_EM_ (brown) gag-specific cells (Fig. 2B, see also Fig 1L-M). A similar pattern was observed across spleen, LNs, BM and blood, with two points to be highlighted (Fig. 2C). First, a statistically significant difference between LNs and blood T_EM_ at d60 (bottom center), not observed at d100 (bottom right), indicating a slow re-entry of LN T_EM_ in G_0_. Second, a tendency of BM T_CM_ cells to contain a smaller fraction of Ki-67^—^ than the other organs, both at d60 (top center) and at d100 (top right), indicating a low level of persistent activation of T_CM_ in the BM. To better investigate the d30-d100 shift, we estimated the absolute numbers of gag-specific cells, and of T_EM_ and T_CM_ gag-specific cells in spleen, LNs, BM and blood (Fig. 2D-F), as well as the absolute numbers of ki 67 ^—^belonging to each subset (Fig. 2G-I). These estimates took into account CD8 T cell abundance in each organ, and the decline of CD8 T cell percentages occurring with aging in spleen, LNs and blood but not in BM (averages: spleen 9.77%; LN 20.85%; blood 9.25%, BM 0.43% at d30, age 11-13 weeks; spleen 7.50%, LNs 15.15%, blood 8.10%, BM 0.50% at d100, age 21-23 weeks). We found that an overall reduction in gag-specific cell numbers coexisted with selective increases of cells belonging to some subsets and/or found in certain organs (Fig. 2D-I). Thus, the sum of gag-specific cells in spleen, LNs, BM and blood at d100 was 88% of that at d30 (Fig. 2D), and that of T_EM_ 76% of that at d30 (Fig. 2F). In striking contrast, taking the four organs altogether, T_CM_ cells increased 2.5 times from d30 to d100 (Fig. 2E), and their percentage among gag-specific cells raised from 7% at d30 to 19% at d100 (Fig. 2D-E). With regards to changes in gag-specific cell distribution, it was remarkable that the sum of cells contained in spleen and blood accounted for ∼ ¾ of the cells found in the four organs altogether at d30, and for only ∼half of them at d100 (Fig. 2D). In fact, from d30 to d100 the number of gag-specific cells was reduced 2.5-fold in blood and a 1.4-fold in spleen, whereas that in LNs and BM increased 1.8- and 1.4-times, respectively (Fig. 2D). Changes of T_EM_ resembled those of gag-specific cells (Fig. 2F), whereas a profound T_CM_ cell increase was observed in all organs, i.e. 2.1-fold in spleen, 2.0-fold in BM, 3.1-fold in LNs, and 3.8-fold in blood (Fig. 2E).

**Figure 2.**
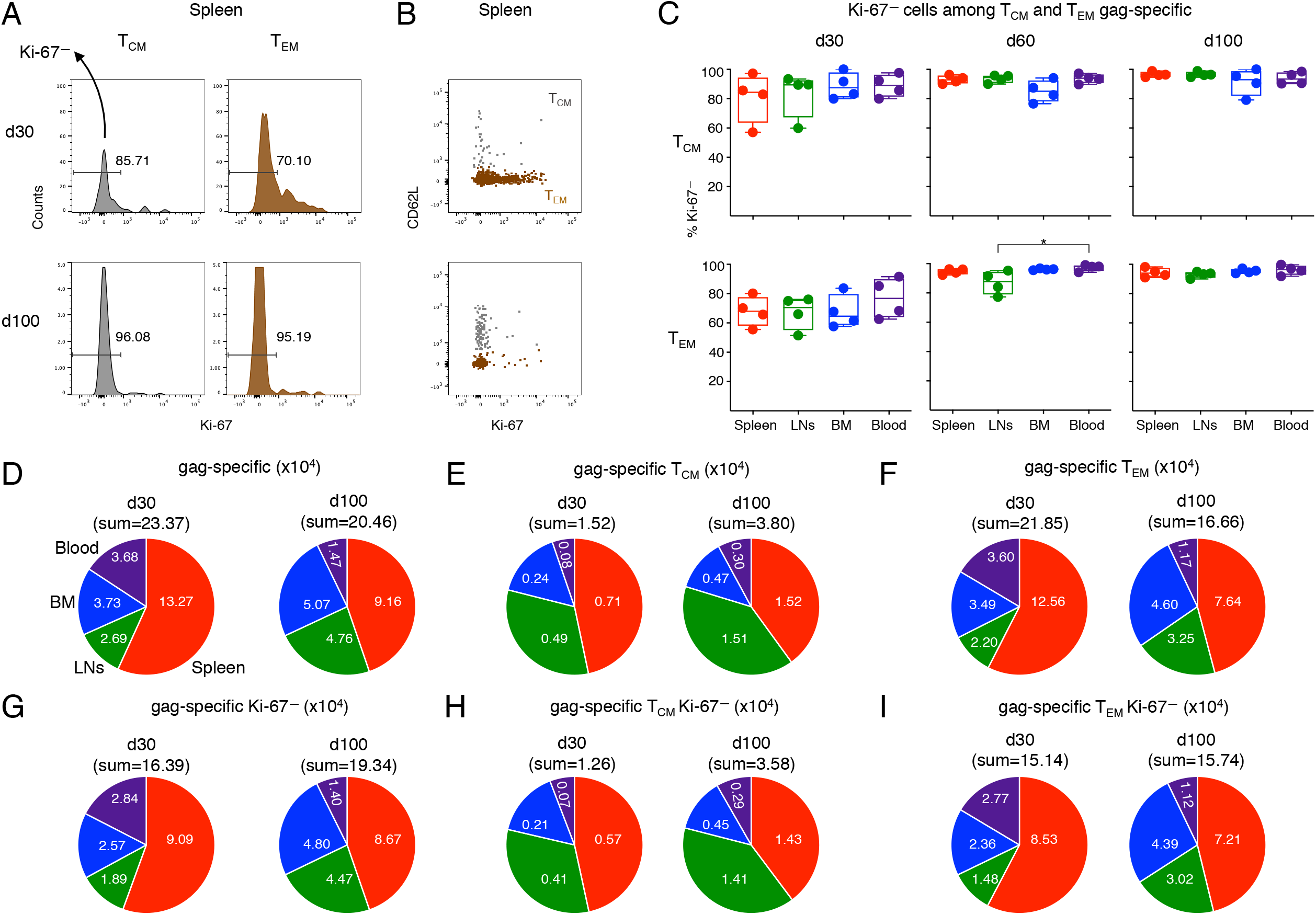
Analysis of ki-67^−^cells among T_CM_ and T_EM_-gag-specific CD8 T cells from ChAd-gag-primed mice, and estimation of absolute cell numbers. Spleen, LN, BM and blood cells from primed mice represented in Fig. 1H-M were analyzed for Ki-67 expression by T_CM_ and T_EM_ gag-specific CD8 T cells, gated as in Fig. 1H and L. **(A-C)**. Typical histograms showing the percentage of ki-67^−^cells among T_CM_ (left) and T_EM_ (right) cells (A), and examples of Ki-67/CD62L plot showing an overlay of T_CM_ (gray) and T_EM_ cells (brown) (B); both panels represent spleen gag-specific CD8 T cells from primed mice at d30 (top) and d100 (bottom). Summary of results of ki-67^−^cells among T_CM_ (top) and T_EM_ (bottom) gag-specific CD8 T cells from spleen, LNs, BM and blood at d30, d60 and d100 (C). In A, the numbers represent the percentages of cells in the indicated regions. **(D-I)**. Absolute numbers of gag-specific (D), T_CM_ (E) and T_EM_ (F) gag-specific CD8 T cells, and of ki-67^−^ cells among each of these subsets (G, H, and I, respectively) at d30 and d100 in spleen, LNs, BM and blood. Statistical analysis was performed by Friedman test with Dunn’s correction for multiple comparison (C). Statistically significant differences are indicated (* *P* ≤ 0.05).

The sum of ki-67^−^ gag-specific cells in spleen, LNs, BM and blood altogether at d100 was 1.2-fold higher than that at d30, as a net result of increase in LNs (2.4-fold) and BM (1.9-fold), almost no change in spleen, and evident reduction in blood (2.0-fold) (Fig. 2G). This was in contrast with the absolute number of ki-67^−^T_CM_ found in the four organs altogether, that showed a striking 2.9-fold increase from d30 to d100, with a pronounced rise in each organ, especially in blood (4.0-fold) and LNs (3.5-fold) (Fig. 2H). The sum of ki-67^−^T_EM_ in the four organs at d100 was similar to that at d30 (Fig. 2I), in contrast to the above-described reduction of total T_EM_ (Fig. 2F). Single organ comparison showed that ki-67^−^T_EM_ cells were reduced in blood (2.5-fold) and spleen (1.2-fold), but increased in LNs (2.0-fold) and BM (1.9-fold) (Fig. 2I).

### RNA sequencing (RNAseq) and bioinformatic analysis of spleen gag-specific CD8 T cells at d30 and d100 after prime

A transcriptomic comparison between d100 and d30 spleen gag-specific cells by RNAseq showed that samples of the two groups were easily separated by Principal Component Analysis (Fig. 3 and S2A). Significant changes at d100 were found in 512 differentially expressed genes (DEGs), 362 of which were down-regulated and 150 up-regulated (Fig 3A). The top 50-up and top 50-down DEGs showed comparable results across samples of each group (Fig. S2B).

**Figure 3.**
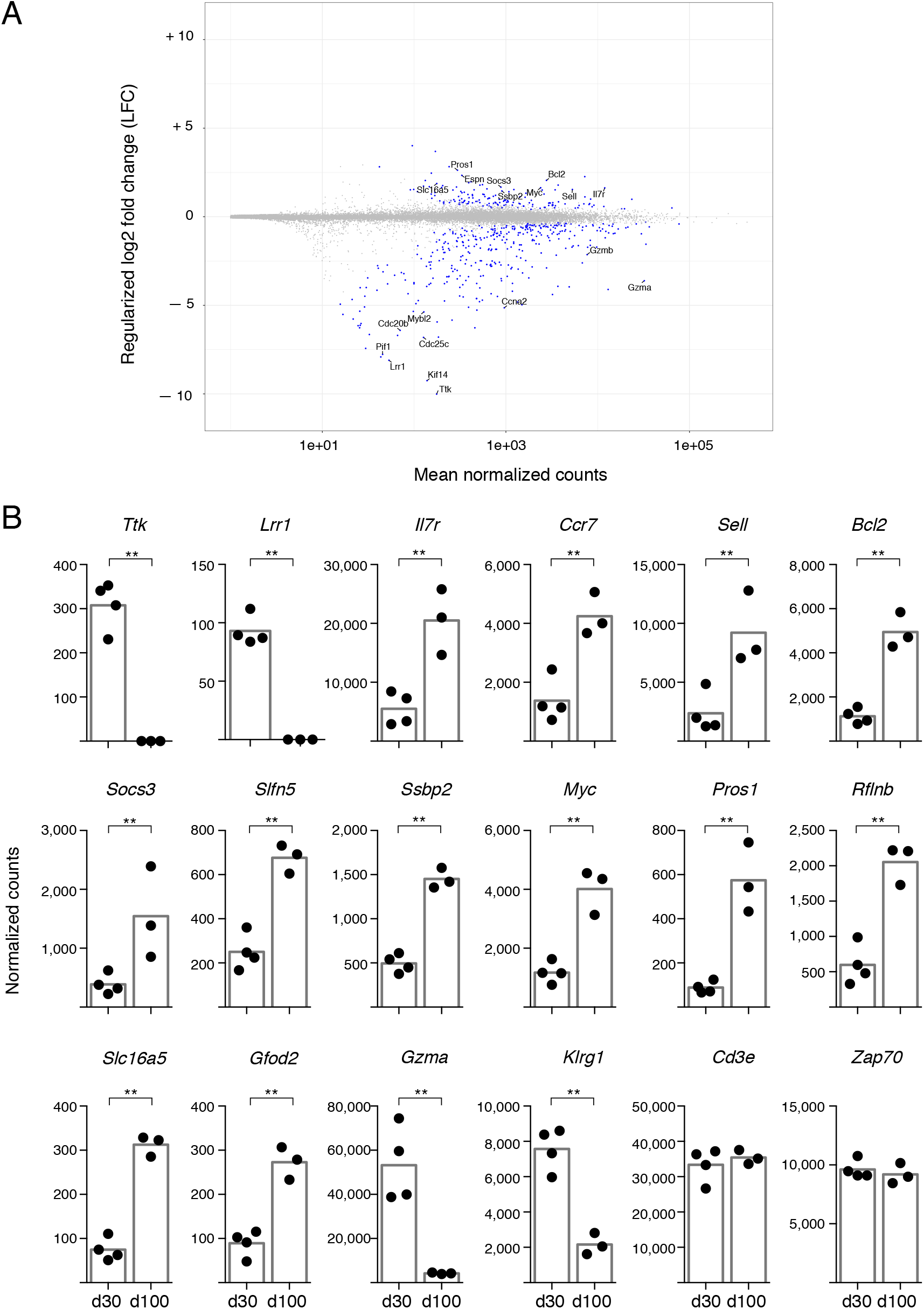
RNA seq analysis of spleen gag-specific CD8 T cells at d30 and d100 post-prime. Bulk RNA was sequenced from sorted spleen gag-specific CD8 T cells at d30 and d100 post-prime, in 4 independent prime experiments with a total of 7 samples. Bioinformatic analysis was performed to identify differentially expressed genes (DEGs) between d100 and d30. **(A)**. Scatterplot of genes whose estimated absolute log-2 fold change was < 11. Genes with a statistically significant difference (false discovery rate (FDR) ≤ 0.01) are highlighted in blue. Log-2 fold changes were regularised using an empirical Bayes method, and mean normalised counts were calculated across all samples. Selected gene symbols among the top 50 significantly upregulated (top 50-up) and the top 50 significantly downregulated (top 50-down) genes are indicated (see Fig. S2 for full list). **(B)**. Normalized counts of representative DEGs having a statistically significant difference between d100 and d30, i.e. *Ttk, Lrr1, IL7r, Ccr7, Sell, Bcl2, Socs3, Slfn5, Ssbp2, Myc, Pros1, Rflnb, Slc16a5, Gfod2, Gzma, Klrg1*. Analysis of *Cd3e* and *Zap70* is included. Each symbol represents an individual sample; columns represent the mean of d30 and d100 group; statistically significant differences are indicated (** FDR ≤ 0.01).

A striking reduction was observed in a small set of downregulated DEGs, with some of them showing a regularized log-2 Fold Change (LFC) comprised between −5 and −10 (Fig 3A). Among the DEGs that were down to virtually 0 normalized counts, *B-myb* (also known as *Mybl2*) was an already recognized T cell effector player of transition into memory state (Powzaniuk et al., 2001; Chen et al., 2018), while others were newly described in this context, e.g. *Ttk*, the gene for Thymidine kinase 1, an IL-2 induced kinase regulating cell cycle progression of T cells (Schmandt et al., 1994); *Kif14*, the gene for Kinesin Family Member 14, a positive regulator of cell cycle (Yang et al., 2019; Zhao et al., 2021); and *Lrr1*, the gene for Leucine-Rich Repeat Protein 1, an inhibitor of 4-1BB signalling (Jang et al., 2001) (Fig. 3A-B). Notably, about half of the top 50 significantly down-regulated genes (top 50-down) encoded for proteins involved in cell proliferation (Fig. S2C, in bold blue).

In contrast, the top 50 up-regulated DEGs (top 50-up) were more heterogeneous, and the LFC did not exceed +5 for any of them (Fig. 3A and Fig. S2C). As expected, the top 50-up DEGs comprised *Ccr7, Sell*, and *Il7r* that encode for CCR7, CD62L, and IL-7Rα, respectively, all recognized markers for T_CM_ phenotype and memory T cell longevity (Kaech et al., 2003; Chen et al., 2018), as well as *Bcl2*, a pro-survival gene (Crauste et al., 2017; Chen et al., 2018), and *Socs3*, which encodes for a cytokine signalling regulator that controls IL-7Rα re-expression after initial down-regulation in activated T cells (Güler et al., 2020) (Fig. 3A-B). Remarkably, top 50-up DEGs included *Slfn5*, a member of the Schlafen family of genes that has been implicated in T cell quiescence and proliferative potential (Geserick et al., 2004; Puck et al., 2017; Metzner et al., 2022), and some genes previously involved in regulating quiescence and maintenance of hematopoietic stem cells (HSCs), i.e. *Ssbp2*, Sequence-specific ssDNA–binding protein 2 (Li et al., 2014), and *Myc* (Wilson et al., 2004), often cited for its role in T cell metabolism and memory T cell differentiation (Marchingo et al., 2020; Nozais et al., 2021; Chen et al., 2018) (Fig. 3A-B). The above mentioned *Socs3* has also been implicated in re-setting quiescence of HSCs after proliferation (Venezia et al., 2004), while the pleiotropic gene *Pros1*, Protein S, also in the top 50-up genes, has been proposed as a regulator of neural stem cell equilibrium between quiescence and proliferation (Zelentsova et al., 2017)(Fig. 3A-B). Altogether, changes in these genes (Fig. 3A-B and Fig. S2C, in bold red) support the notion that the quiescent state of gag-specific cells at d100 was actively and finely regulated.

Additional top 50-up DEGs encoded for proteins regulating cell metabolism and redox state (i.e. *Qpct, Slc16a5, Gfod2, Nmnat3*) (Fig. 3A-B, and Fig. S2C, in bold black), pointing to a metabolic change at d100. It is worth noting that *Rflnb*, Refilin B, a TGF-β effector (Gay et al., 2011) and *Tgfbr3*, TGF-β receptor III, were among the top 50-up DEGs, in agreement with the major role of TGF-β signalling in T cell memory (Ma and Zhang, 2015), and of TGF-β RI and RII in T cell biology (Takai et al., 2013; Ishigame et al., 2013), even though the function of TGF-β RIII has been poorly investigated in this context (Kim et al., 2019). Furthermore, *Cd101* was among the top 50-up DEGs; this gene encodes for a T cell-inhibitory glycoprotein shown to come up in chronic infection (Hudson et al., 2019) (Fig. 3B, and Fig. S2B-C).

Although not listed in the top 50-down, *Gzma, Gzmb*, and *Gzmk* were significantly down-regulated, as were *Nkg7* and *Klrg1*, all typical genes of CD8 T cell effector signature (Fig. 3A-B). Control genes with no significant changes included *CD3d, CD3g, CD3e, ZAP70, Cd8a* and *Cd8b1* (Fig. 3B; RNAseq data available at GSE207389).

### Kinetics of responsiveness to boost

To compare protocols with different prime/boost intervals, ChAd-gag-primed mice were boosted with MVA-gag either at d30 or at d100 post-prime, and rested for 45d. Then the frequency of gag-specific cells, and that of T_CM_ among gag-specific cells, was measured in spleen, LNs, BM and blood (Fig. 4A-B). There was a trend of higher gag-specific frequency when boost was performed at d100 post-prime as compared to boost at d30 in all organs, that reached statistical significance in LNs (Fig. 4A). Independently of the time of boost, the organs with the highest frequencies were BM and blood, followed by spleen and then LNs, similarly to the results at d30 and d60 post-prime (see Fig. 1I). There was no difference between the two boosts in terms of proportion of T_CM_ among gag-specific cells (Fig. 4B). As expected, LNs contained a higher percentage of T_CM_ than any of the other organs, thus resembling post-prime data at d30 and d60 (see Fig. 1M).

**Figure 4.**
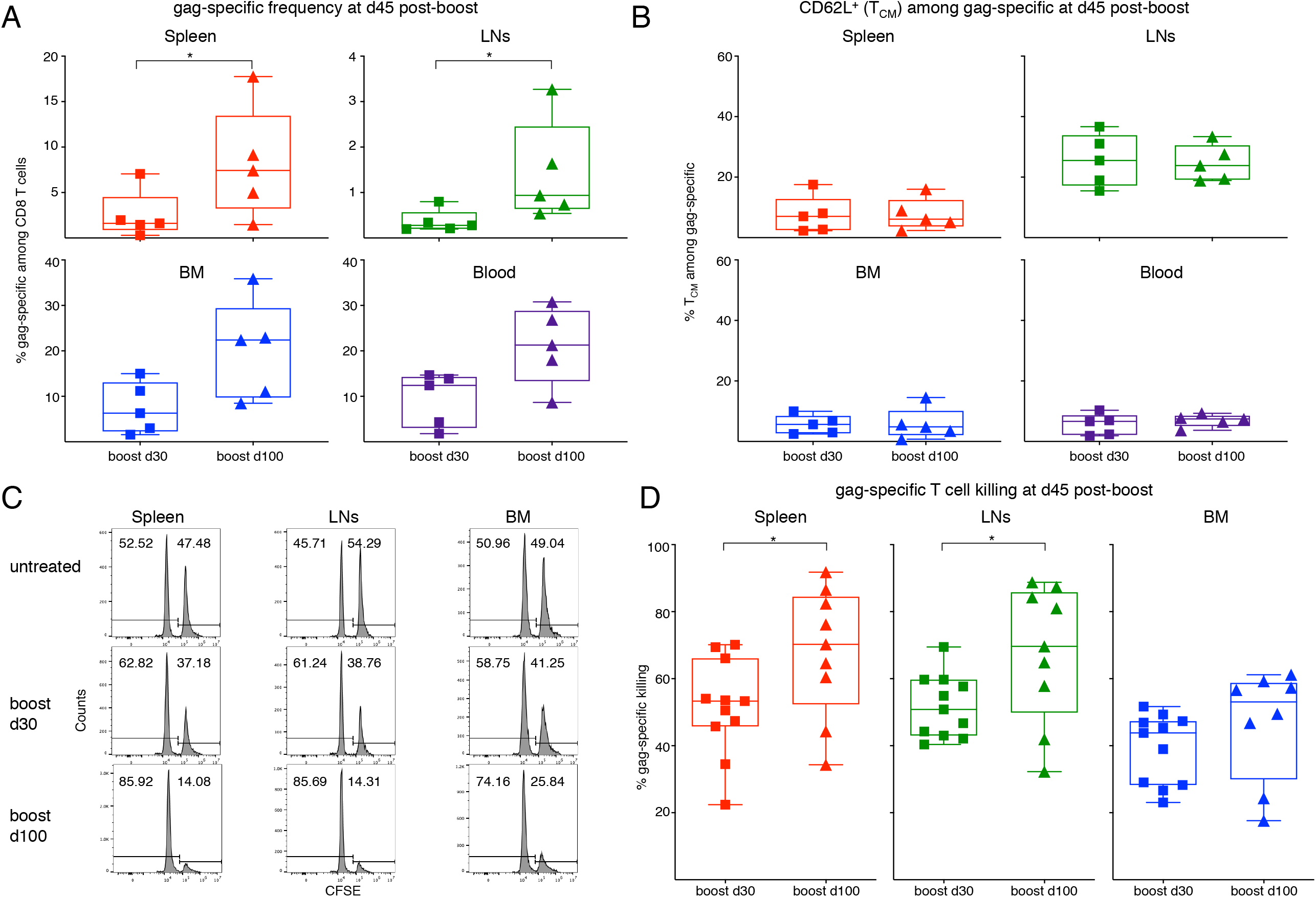
Analysis of gag-specific CD8 T cell frequency, T_CM_-phenotype and *in vivo* killing activity at d45 post-boost. Female BALB/c mice were primed as in Fig. 1 at d0. One set of primed mice was boosted with MVA-gag at d30 post-prime, and another at d100 post-prime. For each set, analysis was performed at d45 post-boost. **(A-B)**. Frequency of gag-specific CD8 T cells (A) and percentage of T_CM_ among gag-specific CD8 T cells (B) in spleen, LNs, BM and blood of primed/boosted mice. **(C-D)**. Primed/boosted and untreated control mice were injected i.v. with a 1:1 mixture of gag-pulsed CFSE^high^ cells and unpulsed CFSE^low^ syngeneic spleen cells (approximately 10×10^6 cells each). After 3 hours, the percentages of CFSE^high^ and CFSE^low^ cells were measured in spleen, LNs and BM, and the percentage of gag-specific killing was determined. Examples of CFSE histograms (C) and summary of results (D). Panels A and B summarize results of 5 independent prime/boost experiments with a total of 60 mice. Each symbol represents a pool of 3 mice. Panel D summarizes results of 4 independent prime/boost experiments with a total of 34 mice. Each symbol represents a single mouse. Statistical analysis was performed by either Student t test, after checking that distribution was normal by Shapiro-Wilk test, or Mann-Whitney test. Statistically significant differences are indicated (* *P* ≤ 0.05

*In vivo* killing assay showed that gag-specific cells were functional in spleen, LNs and BM (Fig. 4C-D). There was a significantly higher percentage of gag-specific killing in spleen and LNs in the group of mice boosted at d100 post-prime as compared to the group boosted at d30, and a tendency of similarly increased killing in the BM (Fig. 4D). Spleen and LNs showed a more prominent killing than BM (Fig. 4C-D), possibly reflecting their higher CD8 T cell percentages (see above). Intracellular IFN-γ assay performed at d45 post-boost by stimulating either spleen or BM cells with a pool of gag protein-derived peptides (gag peptide pool) confirmed that boost at d100 post-prime elicited a stronger CD8 T cell response than boost at d30 (Fig. S3A-C). Moreover, the percentage of IFN-γ^+^ among CD8 T cells was higher in the BM than in the spleen in both experimental groups (Fig. S3B-C), echoing the gag-specific frequency results (see Fig. 4A). Altogether, these results show that boost at d100 was more effective than that at d30.

## Discussion

Most vaccination protocols prescribe administration of at least two vaccine doses, with the aim of achieving sustained protection by first priming and then boosting Ag-specific immunity. Decisions regarding prime/boost time interval have been taken mostly empirically so far (Ledgerwood et al., 2013; Pettini et al., 2021). In this article we described a splenic memory CD8 T cell signature associated with enhanced response to delayed boost in a model of ChAd-gag/MVA-gag vaccination of BALB/c mice.

In our model, delayed boost at d100 post-prime was more effective than early boost at d30 in terms of frequency and phenotype of gag-specific CD8 T cells, *in vivo* gag-specific killing, and IFN-γ production, all measured at d45 post-boost, i.e. in the secondary memory phase. Our data are in the same line of previous evidence showing compromised memory T cell longevity with short 14d-prime/boost intervals (Thompson et al., 2016). However, we specifically addressed memory CD8 T cell maturation, as we focused on ≥ 30 d post-prime, after the ending of clonal expansion (Alvarez-Dominguez and Melton, 2022).

We identified a d100 splenic transcriptomic profile of gag-specific CD8 T cells, characterized by shut off of several proliferative genes (e.g. *Ccna2, Cdc25c, Pclaf*), and up-regulation of stem cell genes previously implicated in setting the equilibrium between quiescence and proliferation (e.g. *Slfn5, Ssbp2, Myc, Socs3*). These results are in agreement with the absolute dominance of ki-67^−^ cells by flow cytometry analysis, and give granularity to the molecular changes occurring from d30 to d100, i.e. a time interval remarkably characterized by stable numbers of ki-67^−^ gag-specific CD8 T cells in the spleen. Transcripts involved in T cell negative regulation and TGF-β pathway (e.g. *Rflnb, Tgfbr3, Cd101*) were also up-regulated, as were some metabolic genes (e.g. *Qpct, Gfod2*). Increased expression of T_CM_/long-lived memory T cell markers (e.g. *Sell, Ccr7, Il7r, Bcl2*), and down-regulation of effector T cell transcripts (e.g. *Gzma, Gzmb, Gzmk, Nkg7, Klrg1*) were in agreement with previous results (Kaech et al., 2002; Chen et al., 2018).

Our flow cytometry analysis and our cell number estimates suggest that gag-specific CD8 T cells migrated from spleen and blood into LNs and BM during the d30-d100 time frame, even though migration in and out of other organs cannot be excluded. Our further findings on gag-specific T_CM_/T_EM_ phenotypes are consistent with the possibility that T_CM_ had a survival advantage over T_EM_ in all examined organs, and likely self-renewed in the BM (Di Rosa, 2016), while a fraction of T_EM_ up-regulated CD62L, thus acquiring a T_CM_ phenotype. Thus, the transcriptomic differences between d100 and d30 spleen gag-specific CD8 T cells might be due to: i) plasticity of the splenic memory population (e.g. T_EM_ to T_CM_ shift); ii) selective survival of a cell subset (e.g. ki-67^−^T_CM_); iii) selective cell recirculation in and out of spleen (e.g. T_EM_ migration out of the spleen and accumulation into LNs and BM); iv) a combination of some or all of the above.

In sum, the main features of the d100 versus d30 gag-specific CD8 T cells in the spleen were a selective 2.6-fold increase in ki-67^−^ T_CM_ as opposed to a 1.2-fold decrease in ki-67^−^T_CM_, and a transcriptional switch to a mature state (Fig. S4). This included, as expected, an increase in T_CM_ phenotype and a decrease in T cell effector genes, plus newly described changes in genes regulating quiescence and proliferation, not implicated before in T cell memory, and in transcripts involved in inhibitory pathways of T cell responses (Fig. S4). It is remarkable that in parallel with the d100-d30 molecular shift in the spleen, we measured significant changes in the blood, i.e. a decline in gag-specific frequency within CD8 T cells, and an increase of T_CM_ within gag-specific cells.

Our data are in agreement with old pioneer studies and more recent ones that altogether emphasize through different approaches the role of a T_CM_ /stem cell molecular profile for long-lived T cell memory (Kaech et al., 2002; Lugli et al., 2013; Graef et al., 2014; Youngblood et al., 2017; Jung et al., 2022), and are consistent with the hypothesis that lymphoid microenvironments regulate the equilibrium between quiescence and self-renewal in long-term T cell memory (Di Rosa, 2016). Notably, our findings establish a new memory CD8 T cell profile of responsiveness to boost, giving a valuable contribution to the rational design of vaccination protocols. Advancements in this field are much needed, as defining the time interval between vaccine shots represents one of the current challenges after the success of many anti-SARS-CoV-2 vaccination strategies, including those based on adenoviral-vectors.

### Limitations of the study

Our study examined female BALB/c mice, which were primed at 7-9 weeks of age. Further research will be required to study mice of both sexes, different ages, and diverse inbred strains. Moreover, we focused our analysis on CD8 T cells from blood, LNs, spleen and BM, and did not examine Tissue Resident Memory CD8 T cells in peripheral tissues, that were reported in other models of vaccination with adenoviral vectors (van der Gracht et al., 2020). Furthermore, our study was limited to ChAd-gag/MVA-gag vaccine model. Additional studies will be required to test different platforms for either homologous or heterologous prime/boost, including mRNA, DNA, etc (Kardani et al., 2016). Finally, we examined Ag-specific CD8 T cells by MHC-I multimers and other assays, but not CD4 T cells.

## Acknowledgements

The following tetramer was obtained through the NIH Tetramer Facility: APC-conjugated H-2K(d) HIV gag 197–205 AMQMLKETI. We thank The Francis Crick Institute’s Advanced Sequencing Facility for RNA sequencing, and Daniel Davies, Silvia Gitto, Magdalene Joseph, Puay Lee, and Alexandru Turcan for useful comments on the manuscript.

Work supported by CTN01_00177_962865 (Medintech) grant from Ministero dell’Università e delle Ricerca (MIUR), by MIUR PRIN grant 2017K55HLC_006, by Reithera, by CNR STM 2021, and by the Francis Crick Institute —which receives its core funding from Cancer Research UK (CRUK), the UK Medical Research Council, and the Wellcome Trust— (FC001003).

## Author contributions

A. Natalini and F. Di Rosa conceived the project, designed experiments, interpreted the results and wrote the paper with help by S. Simonetti. A. Folgori, S. Capone, R. Sbrocchi and A. Nicosia provided the viral vectors and conducted/supervised mouse immunizations; A. Natalini, S. Simonetti G. Favaretto, L. Lucantonio, A.M. Contino performed/analysed flow cytometry experiments; A. Natalini, G. Peruzzi, M. Muñoz-Ruiz, G. Kelly performed/analysed cell sorting and RNAseq experiments; A.M. Contino, R. Sbrocchi, S. Battella performed/analysed intracellular IFN-γ assay. A. Folgori, S. Capone, A. Santoni and A.C. Hayday advised on data discussion and paper writing; A.C. Hayday advised on concepts to prioritize in data analysis and paper writing.

## STAR Methods

### Key resources table

| Reagent or Resource | Source | Identifier |
| --- | --- | --- |
| **Antibodies** |  |  |
| Anti-mouse CD16/CD32 (clone 2.4G2) (Fc block) | BD Biosciences | Cat#: 553142 RRID:AB_394657 |
| PerCP-Cy5.5 anti-mouse CD3ε (clone 145-2C11) | BD Biosciences | Cat#: 551163 RRID:AB_394082 |
| APC anti-mouse CD3ε (clone 145-2C11) | BD Biosciences | Cat#: 553066 RRID:AB_398529 |
| PE anti-mouse CD4 (clone GK1.5) | BD Biosciences | Cat#: 557308 RRID:AB_396634 |
| FITC anti-mouse CD8α (clone clone 53-6.7) | BD Biosciences | Cat#: 553031 RRID:AB_394569 |
| PerCP anti-mouse CD8α (clone 53-6.7) | BD Biosciences | Cat#: 553036 RRID:AB_394573 |
| BUV805 anti-mouse CD8α (clone 53-6.7) | BD Biosciences | Cat#: 564920 RRID:AB_2716856 |
| PE-Cy7 anti-mouse CD62L (clone MEL-14) | BioLegend | Cat#: 104418 RRID:AB_313103 |
| Alexafluor 700 anti-mouse Ki-67 (clone SolA-15) | eBioscience | Cat#: 56-5698-82 RRID:AB_2637480 |
| FITC anti-mouse Ki-67 (clone SolA-15) | eBioscience | Cat#: 11-5698-82 RRID:AB_11151330 |
| Alexafluor 488 anti-mouse CD45 (clone 30-F11) | BioLegend | Cat#: 103121 RRID:AB_1520424 |
| PerCP-Cy5.5 anti-mouse Ter119 (clone TER-119) | eBioscience | Cat# 45-5921-80 RRID:AB_925766 |
| FITC anti-mouse IFN-γ (clone XMG1.2) | BD Biosciences | Cat#: 562019 RRID:AB_10893998 |
| **Chemicals, peptides and recombinant proteins** |  |  |
| gag protein-derived peptides pool (gag peptide pool) | Primm | Customized |
| RPMI 1640 Medium without L-Glutamine with Phenol Red | Euroclone | Cat#: ECB9006L |
| L-Glutamine 100x (200 mM) | Euroclone | Cat#: ECB3000D |
| Penicillin/Streptomycin 100X | Euroclone | Cat#: ECB3001D |
| β-Mercaptoethanol | Sigma | Cat#: M3148 |
| Fetal Bovine Serum | Corning | Cat#: 35-079-CV |
| Trypan blue solution | Euroclone | Cat#: ECM0990D |
| Red blood cell lysis buffer | Sigma | Cat#: R7757 |
| Propidium iodide (PI) | Sigma | Cat#: P4170 |
| RNeasy Plus Mini Kit | Qiagen | Cat#: 74134 |
| Dimethyl sulfoxide (DMSO) | Sigma | Cat#: 276855 |
| Phorbol myristate acetate (PMA) | Sigma | Cat#: P1585 |
| Ionomycin (Iono) | Sigma | Cat#: I0634 |
| Protein Transport Inhibitor<br>(Containing Brefeldin A) (Golgi<br>Plug) | BD Biosciences | Cat#: 555029 |
| **Critical commercial assays** |  |  |
| Fixable Viability Dye eFluor780 | eBioscience | Cat#: 65-0865-14 |
| Fixable Viability Dye Pacific Blue | Invitrogen | Cat#: L23105 |
| Hoechst 33342, Trihydrochloride,<br>Trihydrate | Invitrogen | Cat#: H3570 |
| Carboxyfluorescein succinimidyl<br>ester (CFSE) | eBioscience | Cat#: 65-0850-84 |
| Cytofix/ Cytoperm Kit | BD Biosciences | Cat#: 554714 |
| Foxp3/ Transcription Factor<br>Staining Buffer Set | eBioscience | Cat#: 00-5523-00 |
| **Instruments** |  |  |
| LSRFortessa | BD Biosciences | https://www.bdbiosciences.com/ |
| CytoFLEX B5-R3-V5 | Beckman Coulter | https://www.beckmancoulter.com/ |
| FACSARIA III | BD Biosciences | https://www.bdbiosciences.com/ |
| Illumina HiSeq 4000 | Illumina | https://emea.illumina.com/ |
| **Softwares and algorithms** |  |  |
| FlowJo version 10.7.1 and 10.8.1 | FlowJo LLC | https://www.flowjo.com/ |
| FACS Diva version 6.1.3 | BD Biosciences | https://www.bdbiosciences.com/ |
| Trimmomatic version 0.36 | Usadellab | http://usadellab.org/cms/ |
| R version 4.0.3 |  | https://www.r-project.org/ |
| Prism version 6.0f | GraphPad | https://www.graphpad.com/ |
| **Other** |  |  |
| gag-encoding Chimpanzee<br>Adenovector (ChAd3-gag) | ReiThera | Customized |
| gag-encoding Modified Vaccinia<br>virus Ankara (MVA-gag) | ReiThera | Customized |
| H-2k(d) AMQMLKETI (gag197-205) APC-labeled Tetramer (Tetr-gag) | NIH Tetramer Core<br>Facility | Customized |
| H-2k(d) AMQMLKETI (gag197-205) PE-labeled Pentamer (Pent-gag) | ProImmune | Customized |

## Resource availability

### Lead contact

Further information and requests should be directed to and will be fulfilled by the lead contact, Ambra Natalini.

### Materials availability

This study did not generate new unique reagents.

### Data and code availability

RNAseq data are available in private status: GEO (GSE207389). Data will be publicly available as of the date of publication.

## Experimental model and subjects details

### Adenoviral and MVA vectors

Replication defective, ΔE1 ΔE2 ΔE3 ChAd3 vector encoding HIV-1 gag protein under HCMV promoter (ChAd-gag) and MVA encoding the HIV-1 gag protein under the control of vaccinia p7.5 promoter (MVA-gag) were generated as described and used in all experiments (Colloca et al., 2012; Di Lullo et al., 2009; Di Lullo et al., 2010; Simonetti et al., 2019).

### Mice and vaccination

Six-week-old female BALB/c mice were purchased from Envigo (S. Pietro al Natisone, Udine, Italy), housed at Plaisant animal facility (Castel Romano, Rome, Italy), and used for experiments at 7-9 weeks of age. Mice were divided into groups of at least 35 mice each (untreated and vaccinated). All mice of the vaccinated group were primed at day (d) 0 with ChAd-gag, and either analyzed at the indicated times post-prime, i.e. d30 (range 27-35), d60 (range 60-67), or d100 (range 95-109), or boosted. For prime/boost experiments, primed mice were divided in different sets, each boosted at a single time after prime, i.e. at either d30 or d100 (ranges as above). Viral vectors were administered intramuscularly (i.m.) in the quadriceps at a dose of 10^7^ viral particles (vp) for ChAd-gag and 10^6^ plaque-forming units (pfu) for MVA-gag, in a volume of 50 µl per side (100 µl total) (Simonetti et al., 2019). All experimental procedures were approved by the local animal ethics council and performed in accordance with national and international laws and policies (UE Directive 2010/63/UE; Italian Legislative Decree 26/2014).

## Method details

### Organs

Spleen, LNs, BM and blood were analyzed either at the indicated days after prime or at d45 (range 41-46) after boost. At each time, organs were collected from 3 vaccinated and 3 untreated mice, and cells from the 3 mice of each group were pooled. Spleen, LNs (iliac and inguinal), and blood were processed as described (Simonetti et al., 2019; Quinci et al., 2012). Femurs and tibias were cleared of muscle tissues, and cut at the extremities. The open bone was placed in a cut pipette tip, placed in a microfuge tube, thereby keeping the bone away from the bottom of the tube and allowing the BM to be centrifuged out of the bone at 800x*g* for 1 minute. The bone was discarded and the pellet resuspended, thus obtaining a single cell suspension of BM cells (Abeler-Dörner et al., 2020). All cell suspensions were prepared in RPMI medium with 2 mM L-Glutamine, 100 U/ml Penicillin, 100 µg/ml streptomycin, 50 µM β-Mercaptoethanol + 10% volume/volume (v/v) Fetal Bovine Serum (FBS), and filtered with pre-separation filters (70 µm) (Miltenyi Biotech, Bergisch Gladbach, Germany).

### Cell staining

Cells were incubated with purified anti-mouse CD16/CD32 clone 2.4G2 (Fc block; BD Biosciences, San Jose, CA, USA), and stained as described with H-2k(d) AMQMLKETI (gag_197-205_) allophycocyanin (APC)-labeled Tetramer (Tetr-gag, NIH Tetramer Core Facility, Atlanta, GA, USA) and phycoerythrin (PE)-labeled Pentamer (Pent-gag, Proimmune, Oxford, UK), fluorochrome conjugated monoclonal Antibodies (mAbs) against surface (CD3, CD8 and CD62L) and intracellular (Ki-67) molecules, and Hoechst 33342 (Thermo Fisher Scientific, Waltham, MA, USA) (Simonetti et al., 2019). The following mAbs were used: anti-CD3ε peridinin chlorophyll protein (PerCP)-Cy5.5 (clone 145-2C11, BD Biosciences), anti-CD8α BUV805 (clone 53-6.7, BD Biosciences), anti-CD62L PE-Cy7 (clone MEL-14, Biolegend, San Diego, CA, USA), and anti-Ki-67 mAb conjugated with Fluorescein isothiocyanate (FITC) or Alexafluor 700 (clone SolA-15; eBioscience, Santa Clara, CA, USA). Dead cells were excluded with Fixable Viability Dye conjugated with eFluor780 fluorochrome (eFluor780, Affimetrix, eBioscience)

### Flow cytometry analysis

Samples were analyzed by either LSRFortessa flow cytometer (BD Biosciences) or CytoFLEX System B5-R3-V5 (Beckman Coulter, Brea, CA, USA). In some experiments, CD3^−^ cells were gated out when acquiring spleen and BM samples. Data were analysed using FlowJo software, v. 10.7.1 and 10.8.1 (FlowJo, Ashland, OR, USA).

### *In vivo* killing

Female BALB/c mice were primed with ChAd-gag and boosted with MVA-gag as above. At d45 (range 43-50) post-boost, vaccinated mice and control untreated mice were injected intravenously (i.v.) with 20×10^6 CFSE-labelled spleen cells (from untreated female BALB/c mice), containing a 1:1 mixture of CFSE^high^ gag-peptide pulsed and CFSE^low^ unpulsed cells. After 3 hours, spleen, LNs and BM cells were obtained and analyzed by flow cytometry using a Beckman Coulter Cytoflex instrument. Propidium Iodide (PI) was used for dead cell exclusion. Percentage of gag-specific killing was calculated according to (Barber et al., 2003).

### Cell sorting, RNA sequencing (RNAseq) and bioinformatic analysis

Female BALB/c mice were primed with ChAd-gag as above, and analyzed at either d30 (range 28-29) or d100 (range 96-107) post-prime. CD8 T cells were enriched from pooled RBC-lysed spleen cells of 12 primed mice by negative selection with mouse CD8 Dyna Beads magnetic beads (Thermo Fisher Scientific). Enriched CD8 T cells were stained with Tetr-gag APC, Pent-gag PE, anti-CD3 PerCP-Cy5.5 (as above), anti-CD8α FITC (clone 53-6.7, BD Biosciences) mAbs, and eFluor780. Then live Tetr-gag^+^Pent-gag^+^CD3^+^CD8^+^ cells were sorted by flow cytometry into PBS buffer 1% BSA 2 mM EDTA using a FACSAria III (BD Biosciences) equipped with 488, 561, and 633 nm lasers, and with FACSDiva software (BD Biosciences, v6.1.3). To reduce stress, cells were sorted in gentle FACS-sorting conditions, using a ceramic nozzle of size 100 μm, a low sheet pressure of 19.84 pound-force per square inch (psi) that keeps the sample pressure at 18.96 psi and an acquisition rate of maximum 1500 events/sec. FACS-sorted cells were confirmed to be 92.92 ± 5.70 % pure prior to RNA extraction. Cells were centrifuged for 5 minutes at 300x*g* and pellets resuspended in Buffer RLT Plus (RNeasy Plus Mini Kit from Qiagen, Germantown, MD, USA), and frozen at −80°C. Total RNA was isolated, and cDNA libraries prepared using NEBNext Single Cell/Low Input Library Prep Kit (from 2ng RNA normalised input), following manufacturer’s instructions. They were then sequenced on Illumina HiSeq 4000 with 100bp single-end reads, following manufacturer’s instructions. Read adaptor removal and quality trimming was carried out with Trimmomatic (version 0.36) (Bolger et al., 2014). Reads were then aligned to the mouse genome, using Ensembl GRCm38 - release 95 as reference. Read alignment and gene level quantification was performed by STAR alignment (v.2.5.2a) (Dobin et al., 2013) together with RSEM package (v.1.2.31) (Li and Dewey, 2011). Statistical analyses were performed in the R programming environment (v. 4.0.3). We used DESeq2 (v.1.30.1) to find differential genes using the negative binomial distribution to model counts, with IHW (v.1.18.0) to control the false discovery rate (FDR), and ashr (v. 2.2.47) for effect-size shrinkage. Genes were designated as differentially expressed (DEGs) if FDR ≤ 1.1. All log-fold-changes estimates were regularised using ashr. We used an independent filter of low-signal genes whose effect varied from comparison to comparison, thus the total number of genes was not the same when analyzing down-regulated and up-regulated genes (total number of genes 10,333 and 7,846, respectively) (RNAseq data are available at GEO, access number GSE207389)

### Intracellular IFN-γ assay

Spleen and BM cells were incubated at 37 °C in 5% CO_2_ in round-bottom 96-well plates (2×10^6^ cells/well) for 5 hours with a pool of gag protein-derived peptides (gag peptide pool, 15mers overlapping by 11 amino acids) at final concentration of 2 µg/ml for each peptide. Dimethyl sulfoxide (DMSO, Sigma-Aldrich), the peptide pool diluent, was used as negative control and phorbol myristate acetate/ionomycin (PMA/Iono, Sigma-Aldrich) at final concentration of 20 ng/ml and 1 µg/ml respectively as positive controls. All incubations were performed in the presence of Golgi plug (BD Biosciences). After stimulation, cells were collected and incubated with Fc block, stained with Pacific Blue Live/Dead dye (Invitrogen, Thermo Fisher Scientific) for viability, and with the following mAbs against surface markers: anti-CD3ε APC, clone 145-2C11; anti-CD8α PerCP, clone 53-6.7; anti-CD4 PE, clone GK1.5 (all from BD Biosciences). Intracellular staining was performed after treatment with Fixation/Permeabilization solution and in the presence of Perm/Wash buffer (Cytofix/ Cytoperm kit, BD Biosciences) using anti-mouse IFN-γ FITC, clone XMG1.2 (BD Biosciences).

### Quantification and statistical analysis Estimates of absolute cell numbers

Cells from spleen, LNs and BM were counted by trypan blue exclusion, after lysis of Red Blood Cells (RBCs) (Sigma-Aldrich, St. Louis, MO, USA). Staining with anti-CD45 Alexafluor 488 (clone 30-F11, Biolegend) and anti-Ter119 PerCP-Cy5.5 (clone TER-119, eBioscience) and flow cytometry analysis revealed residual RBCs identified as Ter119^+^ CD45^−^; cells in spleen (on average 40%) and BM (on average 30%), but not in LN samples. Thus, spleen and BM cell counts were multiplied by 0.6 and 0.7, respectively, to obtain nucleated cell counts. Mouse White Blood Cell (WBC) counts/µl and total blood volume were previously reported (Nemzek et al., 2001; Riches et al., 1973). The absolute numbers of gag-specific CD8 T cells in spleen, LNs, BM and blood were estimated based on their percentages determined by flow cytometry and on nucleated cell counts of corresponding organ.

## Statistical analysis

Student t test was used for comparison between two groups whenever each group had ≥9 samples, after checking that distribution was normal by Shapiro-Wilk test. Non-parametric tests were used for the remaining comparisons. Mann-Whitney test was used for comparison between two groups. Either Kruskal-Wallis or Friedman test with Dunn’s correction for multiple comparison were used for comparison among more than two groups. Differences were considered significant when * *P* ≤ 0.05; ** *P* ≤ 0.01. Statistical analysis was performed using Prism v.6.0f, GraphPad Software (La Jolla, CA, USA).

