## Supplementary material for "Improved memory CD8 T cell response to delayed vaccine boost is associated with a distinct molecular signature": 220804-Natalini-Supplementary.pdf

**Supplemental information**

**Supplemental figures**

**Figure S1**

**Analysis of gag-specific CD8 T cells from ChAd-gag-primed mice: gating strategies for flow cytometry analysis, and LN and spleen results.**

**(A). Membrane and Ki-67/DNA-stained sample gating strategy.** Example of gating strategy for flow cytometry analysis of spleen (top) and LN cells (bottom) in 7 steps (gates are indicated in orange, see also ref. (Natalini et al., 2021; Simonetti et al., 2021)): 1) singlet identification based on DNA content; 2) time exclusion, to eliminate any events collected in case of pressure fluctuations; 3) live cell gate; 4) "relaxed" FSC-A/SSC-A (FSC-SSC) gate (the commonly used "narrow" FSC-SSC gate for lymphocytes is shown in white for comparison); 5) CD8 T cell gate; 6) fine exclusion of residual doublets; 7) gag-specific gate, to identify "gag-specific" cells; "not gag-specific" cells were also gated and examined for comparison. Note that we gated out CD3$^{-}$ cells when acquiring spleen samples (step 5, top), and used a stringent criterion for time exclusion to avoid any impact of pressure fluctuations on cell cycle analysis (step 2, bottom). **(B-E). LN results.** LNs from the primed mice represented in Fig. 1A-G were analyzed, and cell cycle of gag-specific (B, top) and not gag-specific (B, bottom) cells was evaluated on DNA/Ki-67 plots as in Fig. 1B. Summary of the kinetics of gag-specific frequency in LNs of primed and untreated mice (C), and of cell cycle phases of gag-specific CD8 T cells in LNs of primed mice (D). Kinetics of the percentages of LN gag-specific CD8 T cells in the "narrow" FSC-SSC gate and in the FSC-A/-H gate (E). **(F). Impact of LN and spleen cell gating on detection of cell cycle phases**. Kinetics of the percentages of LN (left) and spleen (right) gag-specific CD8 T cells in $G_0$, $G_1$, and S-$G_2$/M comprised within either the "narrow" FSC-SSC or the FSC-A/-H gate, as indicated. **(G). Membrane- and Ki-67-stained sample gating strategy.** Example of gating strategy for flow cytometry analysis of spleen cells from untreated (top) and primed (bottom) mice in 6 steps: 1) singlet identification based on FSC-A/-H; 2) time exclusion, to eliminate any events collected in case of pressure fluctuations; 3) live cell gate; 4) "relaxed" FSC-A/SSC-A (FSC-SSC) gate; 5) CD8 T cell gate; 6) gag-specific gate, to identify "gag-specific" cells. A similar strategy was used for LN, BM and blood cell analysis. In flow cytometry plots (in A, B, and G) the numbers represent the percentages of cells in the indicated regions. This figure includes unpublished data in relation to (Simonetti et al., 2019).

**Figure S2. Principal Component Analysis (PCA), and heatmap of the top 50-up and top 50-down statistically significant DEGs.**

Bulk RNAseq and bioinformatic analysis were performed as described in Fig. 3 legend and in materials and methods**. (A).** PCA plot representing samples' variance-stabilised normalised counts projected onto the first two principal components. **(B).** Heatmap of the top 50 significantly upregulated (top 50-up) and of the top 50 significantly downregulated (top 50-down) DEGs, in order of their regularized log2 fold-change estimate (LFC), coloured by variance-stabilised normalised count in a given sample. **(C).** List of the top 50-up and of the top 50-down statistically significant DEGs, ordered as in B. For each DEG the corresponding LFC is indicated. In the top 50-up list, genes regulating quiescence and metabolism are highlighted in bold red and bold black, respectively. In the top 50-down list, genes involved in proliferation (DNA replication, mitosis, cell cycle) are highlighted in bold blue. Please note that *Igkv3-7* expression (LFC -23.56) was found only in 2 out of 4 d30 samples, likely reflecting a B cell contaminant, and in 0 out of 3 d100 samples.

**Figure S3**

**Analysis of intracellular IFN-γ production by gag peptide pool-stimulated CD8 T cells at d45 post-boost.**

Spleen and BM cells from primed/boosted mice from 2 of the 5 experiments represented in Fig. 4A-B were analyzed at d45 post-boost for intracellular IFN-γ production, after stimulation with either gag peptide pool or its diluent DMSO as negative control. In parallel, cells were also stimulated with PMA/Iono as positive control. **(A).** Example of gating strategy for analysis of spleen cells from mice boosted at d30, after incubation with gag peptide pool (top), DMSO (middle), and PMA/Iono (bottom). Live single cells were gated according to the first 3 steps of fig. S1G. Then $CD3^+$ cells were gated on a SSC-A/CD3 plot (left), $CD8^+CD4^-$ cells on a CD4/CD8 plot (center), and IFN-$\gamma^+$ cells on a CD3/IFN-γ plot (right). **(B-C).** Examples of CD3/IFN-γ plots representing spleen (top) and BM cells (bottom) from untreated (left), and primed mice boosted at d30 (center) and at d100 (right) (B), and summary of gag peptide pool-specific IFN-$\gamma^+$ CD8 T cell percentages obtained after subtraction of DMSO background (C). In A and B the numbers represent the percentages of cells in the indicated regions. Panel C summarizes results of 2 independent prime/boost experiments with a total of 18 mice. Statistical analysis was performed by Mann-Whitney test. Statistically significant differences are indicated (* $P \leq 0.05$; ** $P \leq 0.01$).

**Figure S4. Schematic of "quiescent-but highly responsive" memory spleen CD8 T cell signature.**

Spider plot representing key features of splenic signature of gag-specific CD8 T cells at d30 (pink) and d100 (green) post-prime. Units of measure are the following: absolute cell numbers in the spleen for $T_{CM}$, $T_{EM}$, Ki-$67^-$ $T_{CM}$, and Ki-$67^-$ $T_{EM}$ gag-specific CD8 T cells (see Fig. 2E-I); mean normalized counts for *Sell*, *Ssbp2*, *Slnf5*, *Klrg1*, *Ttk*, and *Lrr1* (see Fig. 3B).

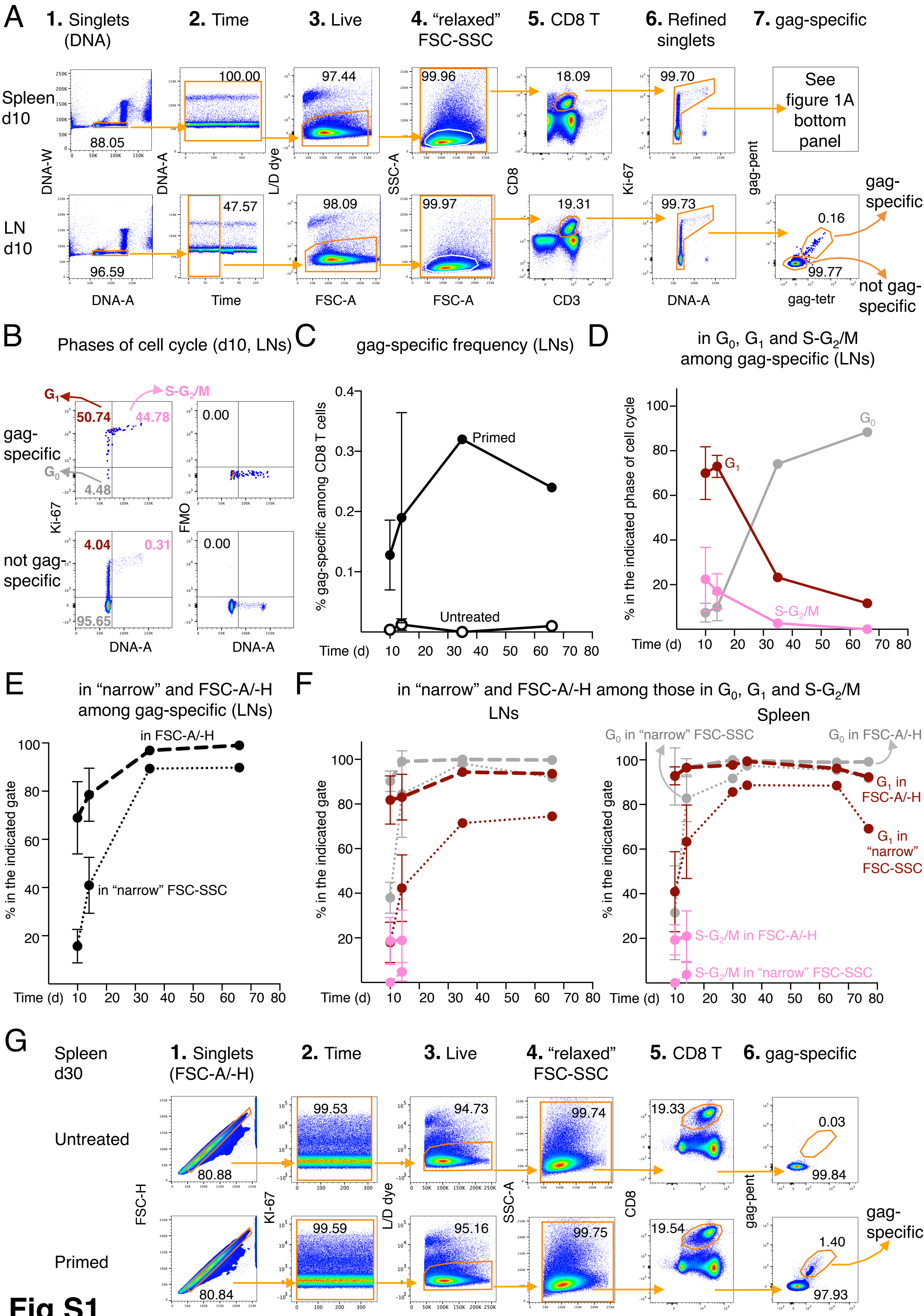
A
1. Singlets (DNA)
2. Time
3. Live
4. "relaxed" FSC-SSC
5. CD8 T
6. Refined singlets
7. gag-specific
Spleen d10
LN d10
88.05
100.00
97.44
99.96
18.09
99.70
See figure 1A bottom panel
96.59
47.57
98.09
99.97
19.31
99.73
0.16
99.77
gag-specific
not gag-specific
DNA-W
DNA-A
L/D dye
SSC-A
CD8
Ki-67
gag-pent
DNA-A
Time
FSC-A
FSC-A
CD3
DNA-A
gag-tetr
B
Phases of cell cycle (d10, LNs)
$G_1$
$S\text{-}G_2/M$
50.74
44.78
$G_0$
4.48
0.00
gag-specific
not gag-specific
4.04
0.31
95.65
0.00
Ki-67
FMO
DNA-A
DNA-A
C
gag-specific frequency (LNs)
% gag-specific among CD8 T cells
Primed
Untreated
Time (d)
D
in $G_0$, $G_1$ and $S\text{-}G_2/M$ among gag-specific (LNs)
% in the indicated phase of cell cycle
$G_0$
$G_1$
$S\text{-}G_2/M$
Time (d)
E
in "narrow" and FSC-A/-H among gag-specific (LNs)
in FSC-A/-H
in "narrow" FSC-SSC
% in the indicated gate
Time (d)
F
in "narrow" and FSC-A/-H among those in $G_0$, $G_1$ and $S\text{-}G_2/M$
LNs
Spleen
$G_0$ in "narrow" FSC-SSC
$G_0$ in FSC-A/-H
$G_1$ in FSC-A/-H
$G_1$ in "narrow" FSC-SSC
$S\text{-}G_2/M$ in FSC-A/-H
$S\text{-}G_2/M$ in "narrow" FSC-SSC
% in the indicated gate
Time (d)
G
Spleen d30
1. Singlets (FSC-A/-H)
2. Time
3. Live
4. "relaxed" FSC-SSC
5. CD8 T
6. gag-specific
Untreated
Primed
80.88
99.53
94.73
99.74
19.33
0.03
99.84
80.84
99.59
95.16
99.75
19.54
1.40
97.93
gag-specific
FSC-H
KI-67
L/D dye
SSC-A
CD8
gag-pent
FSC-A
Time
FSC-A
FSC-A
CD3
gag-tetr


Fig S1

### Fig. S2

A

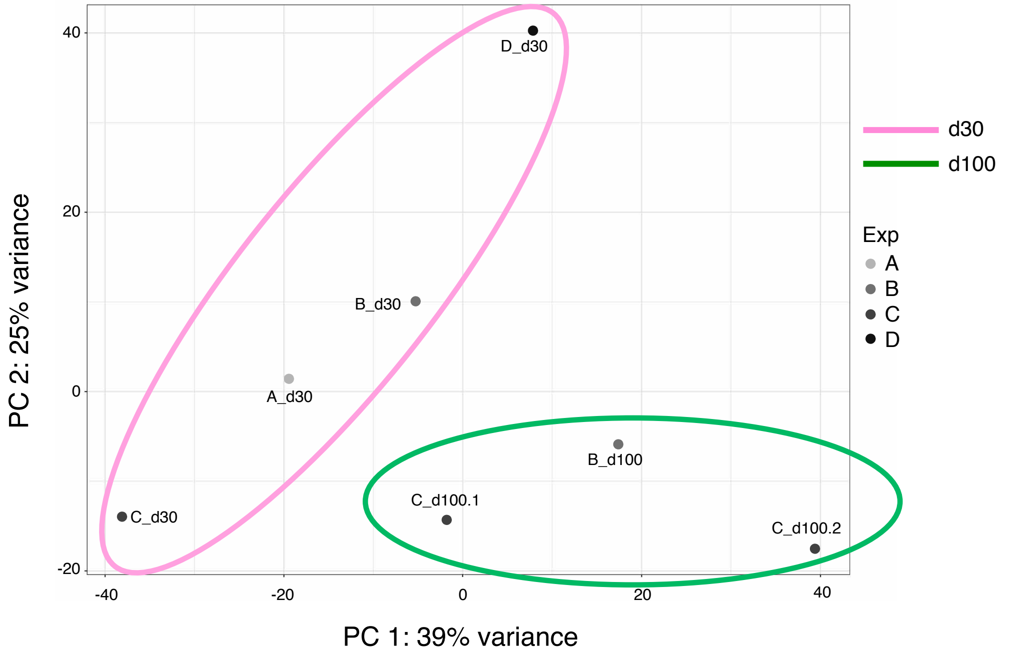


B

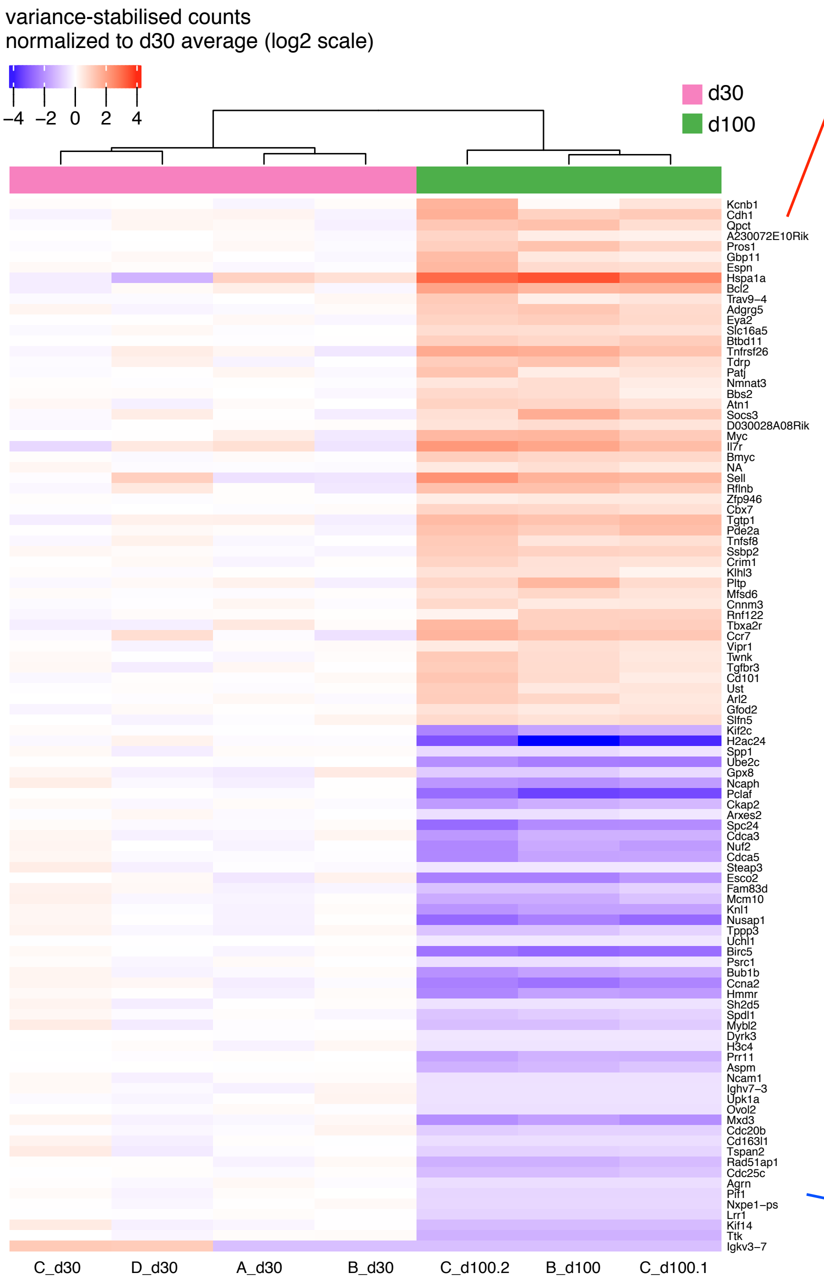


C

| TOP 50-UP | LFC |
|---|---|
| *Kcnb1* | 4.01 |
| *Cdh1* | 3.69 |
| ***Qpct*** | **2.83** |
| *A230072E10Rik* | 2.82 |
| ***Pros1*** | **2.63** |
| *Gbp11* | 2.46 |
| *Espn* | 2.34 |
| *Hspa1a* | 2.26 |
| *Bcl2* | 2.05 |
| *Trav9-4* | 2.05 |
| *Adgrg5* | 1.98 |
| *Eya2* | 1.92 |
| ***Slc16a5*** | **1.87** |
| *Btbd11* | 1.81 |
| *Tnfrsf26* | 1.79 |
| *Tdrp* | 1.78 |
| *Patj* | 1.76 |
| ***Nmnat3*** | **1.73** |
| *Bbs2* | 1.70 |
| *Atn1* | 1.68 |
| ***Socs3*** | **1.67** |
| *D030028A08Rik* | 1.65 |
| ***Myc*** | **1.64** |
| *Il7r* | 1.63 |
| *Bmyc* | 1.57 |
| *ENSMUSG00000116180* | 1.54 |
| *Sell* | 1.54 |
| *Rflnb* | 1.53 |
| *Zfp946* | 1.52 |
| *Cbx7* | 1.52 |
| *Tgtp1* | 1.52 |
| *Pde2a* | 1.51 |
| *Tnfsf8* | 1.51 |
| ***Ssbp2*** | **1.45** |
| *Crim1* | 1.43 |
| *Klhl3* | 1.42 |
| *Pltp* | 1.41 |
| *Mfsd6* | 1.38 |
| *Cnnm3* | 1.37 |
| *Rnf122* | 1.35 |
| *Tbxa2r* | 1.33 |
| *Ccr7* | 1.28 |
| *Vipr1* | 1.28 |
| *Twnk* | 1.27 |
| *Tgfbr3* | 1.25 |
| *Cd101* | 1.25 |
| *Ust* | 1.25 |
| *Arl2* | 1.21 |
| ***Gfod2*** | **1.20** |
| ***Slfn5*** | **1.19** |

| TOP 50-DOWN | LFC |
|---|---|
| ***Kif2c*** | **-4.02** |
| *H2ac24* | -4.11 |
| *Spp1* | -4.15 |
| *Ube2c* | -4.21 |
| *Gpx8* | -4.35 |
| ***Ncaph*** | **-4.35** |
| ***Pclaf*** | **-4.39** |
| *Ckap2* | -4.47 |
| *Arxes2* | -4.49 |
| ***Spc24*** | **-4.50** |
| ***Cdca3*** | **-4.53** |
| ***Nuf2*** | **-4.62** |
| ***Cdca5*** | **-4.62** |
| *Steap3* | -4.67 |
| ***Esco2*** | **-4.71** |
| ***Fam83d*** | **-4.73** |
| ***Mcm10*** | **-4.75** |
| ***Knl1*** | **-4.80** |
| ***Nusap1*** | **-4.82** |
| *Tppp3* | -4.91 |
| *Uchl1* | -4.94 |
| *Birc5* | -4.97 |
| ***Psrc1*** | **-5.09** |
| ***Bub1b*** | **-5.11** |
| ***Ccna2*** | **-5.14** |
| *Hmmr* | -5.27 |
| *Sh2d5* | -5.33 |
| ***Spdl1*** | **-5.35** |
| *Mybl2* | -5.37 |
| *Dyrk3* | -5.51 |
| *H3c4* | -5.77 |
| ***Prr11*** | **-5.84** |
| ***Aspm*** | **-5.94** |
| *Ncam1* | -6.03 |
| *Ighv7-3* | -6.13 |
| *Upk1a* | -6.15 |
| *Ovol2* | -6.27 |
| *Mxd3* | -6.27 |
| ***Cdc20b*** | **-6.41** |
| *Cd163l1* | -6.65 |
| *Tspan2* | -6.70 |
| *Rad51ap1* | -6.80 |
| ***Cdc25c*** | **-6.81** |
| *Agrn* | -7.43 |
| *Pif1* | -7.77 |
| *Nxpe1-ps* | -7.91 |
| *Lrr1* | -8.10 |
| ***Kif14*** | **-9.25** |
| ***Ttk*** | **-10.01** |
| *Igkv3-7* | -23.56 |

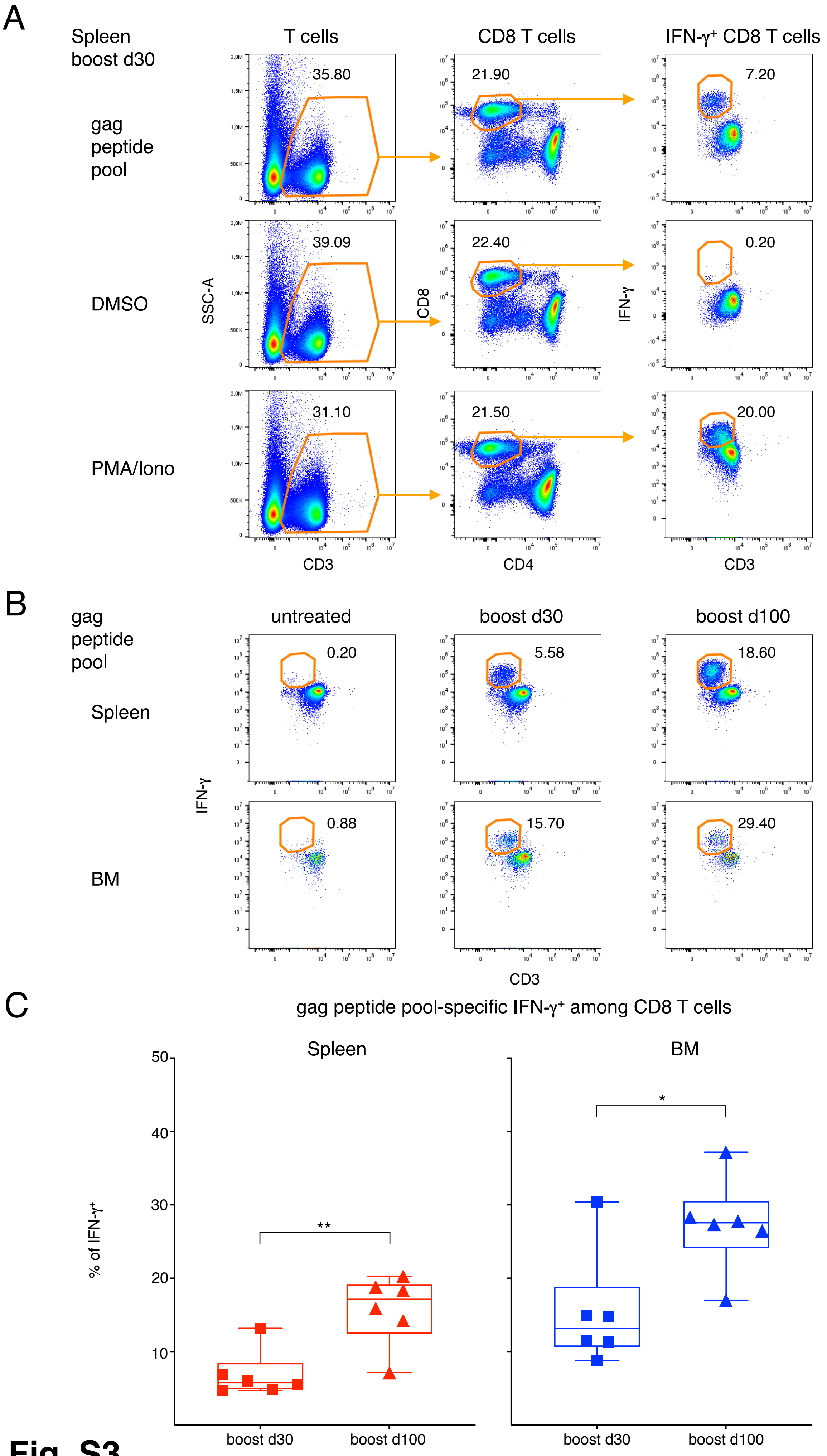
A
Spleen
boost d30
T cells
CD8 T cells
IFN-γ⁺ CD8 T cells
gag
peptide
pool
DMSO
PMA/Iono
35.80
21.90
7.20
39.09
22.40
0.20
31.10
21.50
20.00
SSC-A
CD8
IFN-γ
CD3
CD4
CD3
B
gag
peptide
pool
untreated
boost d30
boost d100
Spleen
BM
0.20
5.58
18.60
0.88
15.70
29.40
IFN-γ
CD3
C
gag peptide pool-specific IFN-γ⁺ among CD8 T cells
Spleen
BM
% of IFN-γ⁺
**
*
boost d30
boost d100
boost d30
boost d100

**Fig. S3**

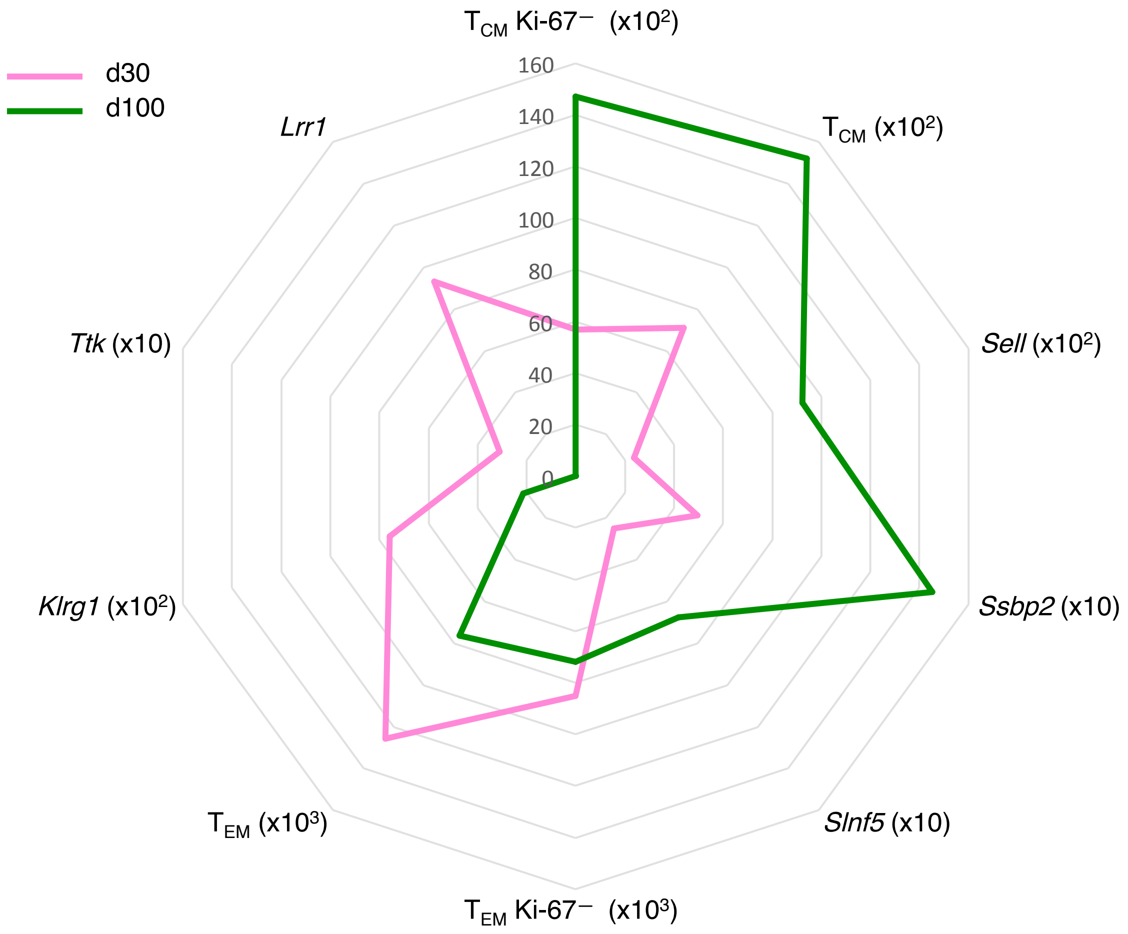
d30
d100
T CM Ki-67⁻ (x10²)
T CM (x10²)
Sell (x10²)
Ssbp2 (x10)
Slnf5 (x10)
T EM Ki-67⁻ (x10³)
T EM (x10³)
Klrg1 (x10²)
Ttk (x10)
Lrr1
160
140
120
100
80
60
40
20
0

**Fig. S4**
